# Periostin facilitates ovarian cancer recurrence by enhancing cancer stemness

**DOI:** 10.1101/2023.03.30.534465

**Authors:** Zhiqing Huang, Olivia Byrd, Sarah Tan, Bailey Knight, Gaomong Lo, Lila Taylor, Andrew Berchuck, Susan K. Murphy

**Affiliations:** Duke University School of Medicine, Department of Obstetrics and Gynecology, Division of Reproductive Sciences; Duke University School of Medicine, Department of Obstetrics and Gynecology, Division of Gynecologic Oncology

**Keywords:** Primary ovarian cancer (pOC), recurrent ovarian cancer (rOC), Periostin (POSTN), tumor microenvironment (TME), extracellular matrix (ECM), chemotherapeutic response, conditioned medium (CM)

## Abstract

Ovarian cancer (OC) is the deadliest reproductive system cancer. Its high lethality is due to the high recurrence rate and the development of chemotherapeutic resistance, which requires synergy between cancer cells and non-cancerous cells of the tumor microenvironment (TME). Analysis of gene expression microarray data from paired primary and recurrent OC tissues revealed significantly elevated expression of the gene encoding periostin (*POSTN*) in recurrent OC compared to matched primary tumors (p=0.014). Finding POSTN primarily localized to the TME, we investigated the role of TME POSTN in OC cell viability, migration/invasion, and chemosensitivity. Conditioned media with high levels of POSTN (CM*^POSTNhigh^*) was generated using *POSTN*-transfected fibroblastic preadipocyte 3T3-L1 cells. CM*^POSTNhigh^*-cultured OC cells exhibited faster migration, more invasiveness (p=0.006), and more chemoresistance (p<0.05) compared to OC cells cultured with control medium (CM*^CTL^*). Furthermore, CM*^POSTNhigh^*-cultured HEYA8 cells demonstrated increased resistance to paxlitaxel-induced apoptosis. Multiple OC cell lines (HEYA8, CAOV2, and SKOV3) cultured with CM*^POSTNhigh^*showed increases in stem cell side population relative to CM*^CTL^*-cultured cells. *POSTN*-transfected 3T3-L1 cells exhibited more intracellular and extracellular lipids, and this was linked to increased cancer cell expression of the oncogene fatty acid synthetase (FASN). Additionally, POSTN functions in the TME were linked to Akt pathway activities. In a xenograft mouse model of OC, the mean tumor volume in mice injected with CM*^POSTNhigh^*-grown OC cells was larger than that in mice injected with CM*^CTL^*-grown OC cells (p=0.0023). Altogether, higher *POSTN* expression is present in recurrent OC and promotes a more aggressive and chemoresistant oncogenic phenotype *in vitro.* Within cancer TME fibroblasts, POSTN can stimulate lipid production and is associated with increased OC stem cell side population, consistent with its known role in maintaining stemness. Our results bolster the need for further study of POSTN as a potential therapeutic target in treatment and potential prevention of recurrent ovarian cancer.

**Author Summary:** Ovarian cancer has a high rate of recurrent disease that is often resistant to chemotherapy. Comparing primary and recurrent ovarian cancer tumors, we found that the gene *POSTN*, which encodes the protein periostin, is more highly expressed in recurrent tumors, and more highly expressed in the tumor microenvironment, outside of the cancer cells. We transfected cells with vectors encoding POSTN or with blank vectors to generate conditioned media with high POSTN or control media. Ovarian cancer cells cultured in the POSTN-high conditioned media showed faster wound healing, more invasiveness, and more resistance to apoptosis caused by chemotherapeutic agents, and increased stemness, an important trait in cancer cells, especially recurrent cells. POSTN-transfected cells showed higher expression of the enzyme fatty acid synthase and higher concentrations of lipids, indicating that POSTN may play a role in increasing the energy available to cancer cells. The Akt pathway, often activated in ovarian cancer growth, was activated more in cells cultured in the POSTN-high environment. Finally, we injected immunocompromised mice with ovarian cancer cells that were grown in either the POSTN-high media or the control media, and the average tumor size was higher in mice injected with the cells that were grown in the POSTN-high media.

## Introduction

In the United States during the year 2023, there will be approximately 19,710 new cases of ovarian cancer (OC) and 13,270 deaths due to ovarian cancer ^1^. OC is the second-most-common gynecologic malignancy, accounting for more deaths than any other cancer of the female reproductive system ^2^. The main reasons for the higher death rate associated with ovarian cancer are that it often goes undetected or misdiagnosed due to a lack of specific symptoms, the frequency of OC recurrence, and intrinsic or acquired chemoresistance. Treatment strategies have not significantly improved in the past 30 years, and immunotherapy is not effective with most OC patients. Currently, most patients get neoadjuvant chemotherapy followed by interval debulking and then finish chemotherapy ^3^. About half of the patients get maintenance treatment with PARP inhibitors ^4^. Yet 70-75% of individuals diagnosed with advanced stage serous OC will experience recurrent, incurable disease despite an initial promising response to treatment ^5–7^. Although the genetics of primary OC have been extensively studied, little data is available on recurrent tumors as it is difficult to obtain recurrent tumor samples and even more difficult to obtain primary (pOC) – recurrent (rOC) tumor pairs ^7^. Investigation into the genomic differences that characterize rOC, and how these differences contribute to tumor progression and chemotherapeutic response is of great importance in the effort to identify relevant targets that allow for more effective treatment of patients with OC.

Tumor heterogeneity and the need to minimize toxicity to normal cells presents a tremendous challenge for treating chemoresistant disease ^8^. The tumor microenvironment (TME) plays an important role in the development of acquired chemoresistance ^9, 10^. The TME includes the blood vessels, immune cells, fibroblasts, signaling molecules, and extracellular matrix (ECM) that surround a tumor ^11, 12^. The ECM is perturbed in tumors and can promote the growth, survival, and invasion of cancer; the ECM also modifies fibroblast and immune cell behavior to promote metastasis and impair response to treatment ^13^. The ECM is a complex assembly of fibrous proteins, proteoglycans, and other molecules, including cytokines and growth factors ^14^. Matricellular proteins play a central role in the homeostasis of normal tissues regulating cell proliferation and differentiation ^14, 15^. These proteins are generally expressed at low levels in most adult tissues but are highly expressed during inflammation, tissue repair, wound healing, and malignant transformation ^15^. Among these, Periostin (*POSTN*), also termed osteoblast-specific factor 2, is a key player in tissue repair and remodeling and plays a role in several immune-mediated inflammatory conditions and in cancer development and progression ^16^. The *POSTN* gene encodes a secreted extracellular matrix protein, Periostin, that is required for maintaining the cell microenvironment during normal cell growth and proliferation. *POSTN*-encoded protein binds to integrins to support adhesion and migration of epithelial cells ^17^. Elevated expressions of the *POSTN* gene and Periostin protein have been reported in cancer cells and in the TME ^16^ and is associated with poor prognosis as well as resistance to chemotherapeutic treatments, including in OC ^18^.

POSTN is known to be involved in the promotion of cancer cell growth, cancer invasion, and chemoresistance ^16^, but the role of POSTN in the TME, especially concerning its contribution to cancer recurrence, has not been fully investigated. Given the very strong relationship between recurrent OC and mortality, we focused our investigation on genomic differences between primary and recurrent OC, intending to exploit these differences for the development of novel targeted therapies. This study examined the role of exogenous *POSTN* in OC proliferation, migration/invasion, and chemosensitivity, and the regulatory POSTN pathway as a component of the TME in ovarian cancer.

## Results

### Periostin is highly expressed in recurrent ovarian cancer

To investigate gene expression changes in tumors that distinguish recurrent from primary ovarian cancers, we generated gene expression microarray data using Affymetrix U133 Plus 2.0 arrays for 16 matched primary (pOC) and recurrent high grade serous epithelial ovarian cancer tissues (rOC) ^19, 20^. RMA normalized gene expression values were compared between these two groups using paired student t-tests. There were 642 genes showing significant differential expression between pOCs and rOCs (p<0.05). Among these genes, nine exhibited a greater than two-fold difference in expression (**Figure 1A**). Six of these had more than two-fold higher expression in rOC, including periostin (*POSTN*), collagen type XI alpha 1 chain (*COL11A1*), tenascin C (*TNC*), asporin (*ASPN*), matrix metalloproteinases 13 (*MMP13*), and leucine rich repeat containing 15 (*LRRC15*). Three exhibited more than two-fold lower expression in rOC, including complement C7 (*C7*), paternally expressed gene 3 (*PEG3*), and tetraspanin 8 (*TSPAN8*). Interestingly, all six genes with elevated expression in rOC vs pOC belong to the extracellular matrix (ECM) family. Of these, *POSTN* showed the most marked difference in expression between the groups, with 11 of the 16 pairs exhibiting higher expression in rOC (RMA expression values: pOC, 7.73; rOC, 10.23; p=0.014) (**Figure 1B**).

**Figure 1.**
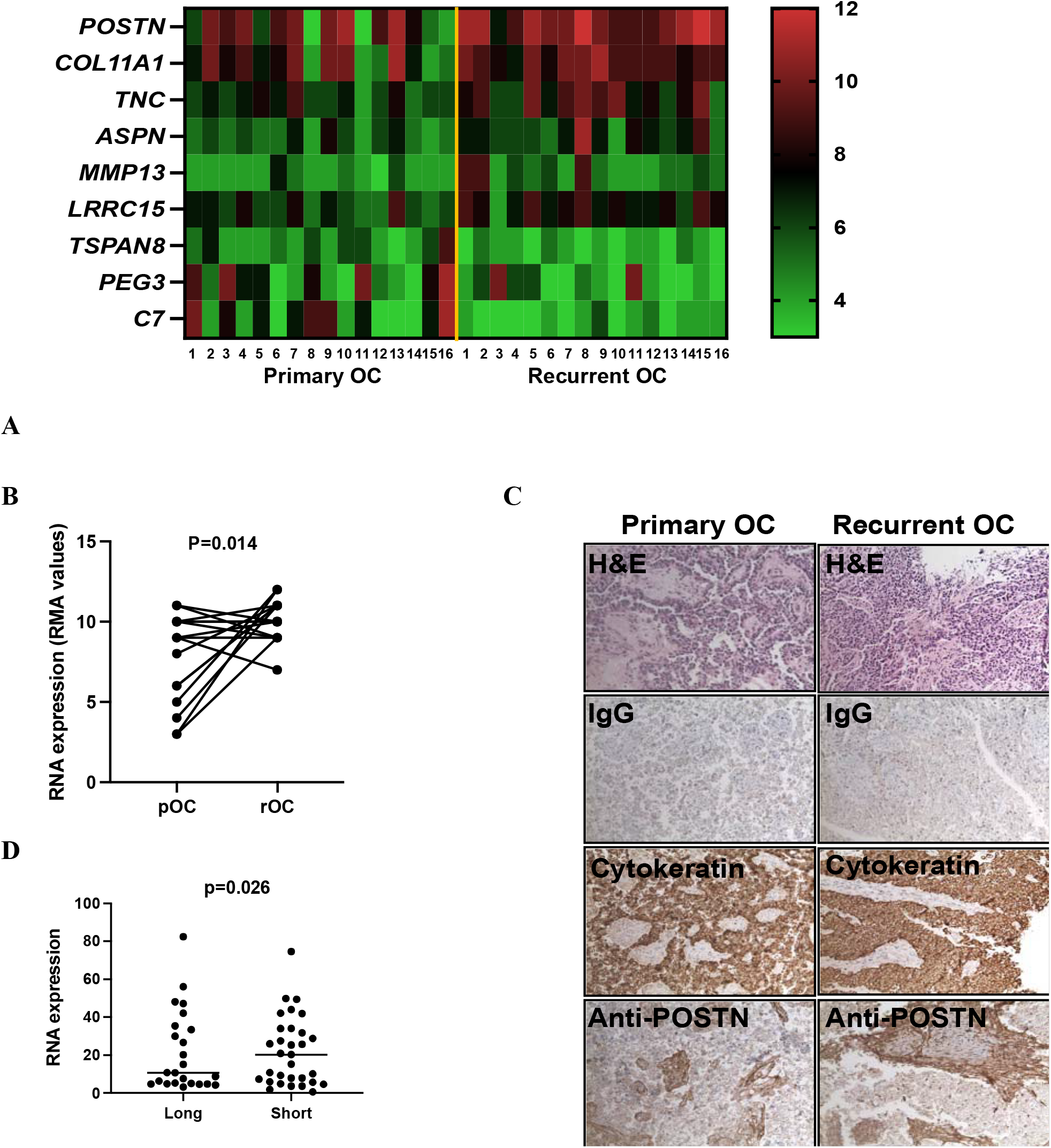
Periostin is highly expressed in recurrent ovarian cancer and is associated with worse prognosis for ovarian cancer patients. A. Gene expression microarray results for sixteen paired pOC and rOC tissues. Shown are data from 9 genes with >2-fold differences in expression. B. *POSTN* expression in each pOC and rOC pair. *POSTN* was increased in the rOC specimen for 11 of the 16 pairs (p=0.014). C. Elevated POSTN protein was evident in rOC relative to pOC with higher levels present in the stroma. Micrographs were taken at total magnification of 100x. D. *POSTN* expression in high grade serous OC from women with <3 years (short) vs >7 years (long) survival (p=0.026).

Immunohistochemistry staining confirmed that, like the mRNA levels, POSTN protein levels were also elevated in rOC (**Figure 1C**) relative to the matched pOC and showed higher levels in stromal tissues than in the cancer cells.

We next examined *POSTN* expression in an independent dataset of high-grade serous ovarian cancer run on the same array platform as our pOC-rOC paired samples ^21^, including 31 tumors from women who lived more than seven years post-diagnosis and 24 tumors from women who lived less than three years post-diagnosis (**Figure 1D**). In this dataset, *POSTN* exhibited higher expression in the short-term survivors than in the long-term survivors (p=0.026) (**Figure 1D**). In summary, *POSTN* expression is higher in rOC than in pOC and in tumors from patients with shorter-term rather than longer-term survival post-diagnosis, with protein expression most prominent in cancer stromal tissues. *POSTN* expression is inversely associated with ovarian cancer prognosis.

### Extracellular POSTN enhances invasiveness and chemoresistance in vitro

Consistent with prior reports ^22–24^, we found that cancer stromal tissues express high levels of *POSTN* gene and the level of stromal *POSTN* is inversely related to prognosis (**Figure 1C and 1D**). However, the effects of elevated exogenous levels of POSTN in cancers, such as in TME tissues, have not been extensively investigated. We generated stable mouse 3T3-L1 preadipocyte cell lines with plasmid pLenti-GIII-CMV-RFP-2A-Puro-*POSTN* that results in high levels of POSTN protein expression, and a control plasmid (pLenti-CMV-RFP-2A-Puro-Blank Vector).

Due to the co-expression of red fluorescent protein (RFP) in cells, we were able to select *POSTN+* cells using fluorescence-activated cell sorting with the RFP marker. RT-PCR with a *POSTN* probe showed significantly higher expression of *POSTN* in 3T3-L1 cells (**Figure 2A**, left panel, p=0.003). In **Figure 2A** (right), ELISA was performed using the medium collected from either *POSTN*-transfected 3T3-L1 cells (3T3-*POSTN*) or vector-transfected cells (3T3-Blank) following at least 48 hours in cell culture. The conditioned medium (CM) from 3T3-*POSTN* cells showed a significant induction of POSTN protein (CM*^POSTNhigh^*) compared to CM from 3T3-blank cells (CM*^CTL^*, p=0.02).

**Figure 2.**
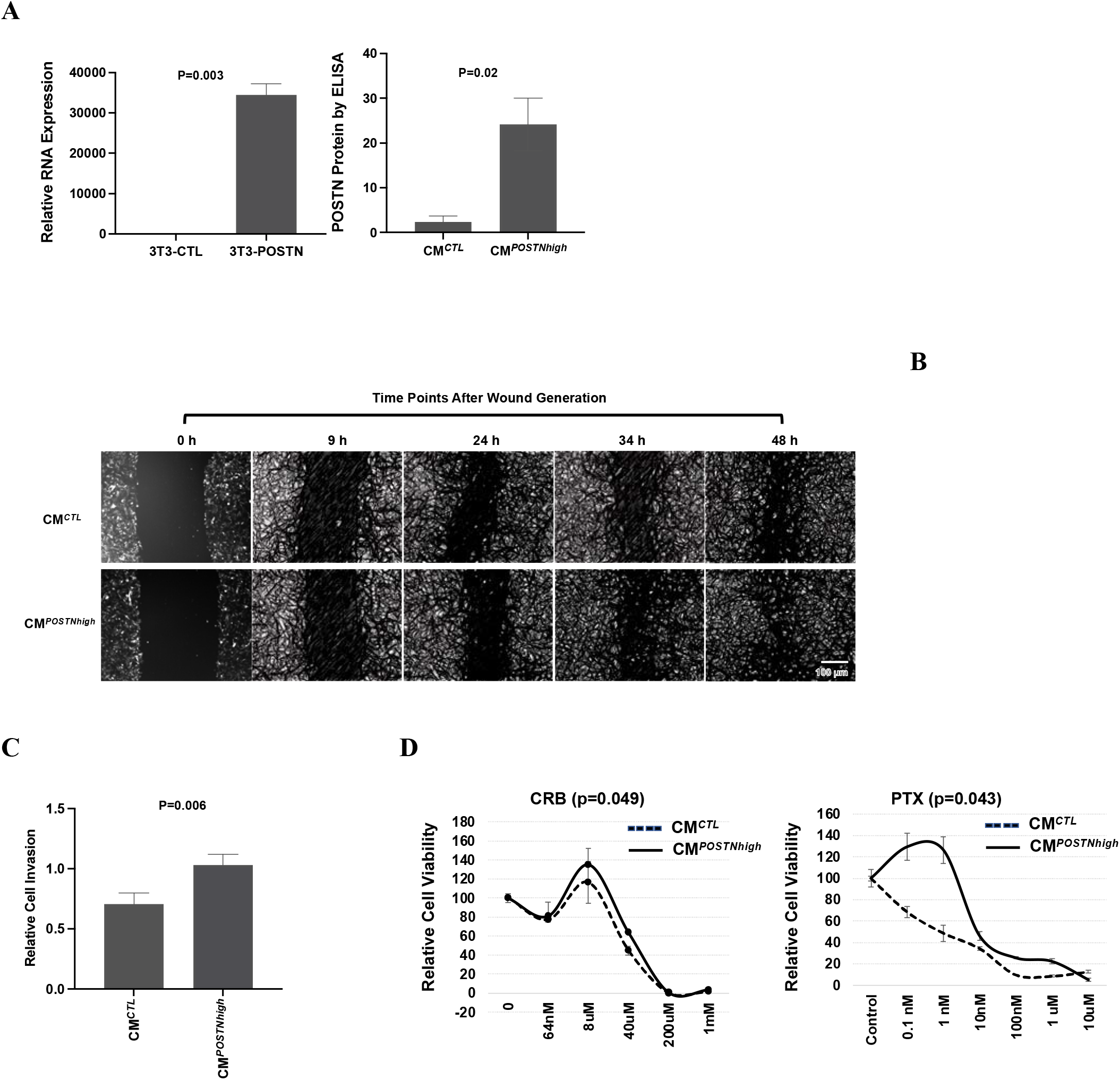
Higher *POSTN* expression in the tumor environment leads to more aggressive cancer phenotypes. **A.** Generation of conditioned medium in 3T3-L1 cells with high levels of POSTN. Left, *POSTN* RNA expression in 3T3 cells stably transfected with a *POSTN* expression vector (3T3-POSTN) relative to the blank vector infected cells (3T3-CTL), normalized with the internal control β2M (p=0.003). Right, POSTN protein levels in the conditioned medium from 3T3-POSTN cells (CM*^POSTNhigh^*) compared to 3T3-CTL cells (CM*^CTL^*), normalized to total protein levers in cells (p=0.02). **B.** HEYA8 cells exhibit faster migration and proliferation (‘wound healing’) under CM*^POSTNhigh^* than CM*^CTL^*. (Scale bar, 100 µm; 4x magnification). **C.** HEYA8 cells are more invasive when cultured under CM*^POSTNhigh^* vs CM*^CTL^*(p=0.006). **D.** Culture under CM*^POSTNhigh^* vs CM*^CTL^*promotes chemoresistance in A2780 cells (p=0.049 for carboplatin [CRB] and 0.043 for paclitaxel [PTX].

To determine the impact of POSTN on cell migration, the ovarian cancer cell line HEYA8 was cultured in media containing CM*^POSTNhigh^* or CM*^CTL^* and assessed with a wound healing assay. As shown in **Figure 2B**, HEYA8 cells cultured with CM*^POSTNhigh^* exhibited faster cell migration, evidenced by the shorter time taken to close the gap that was introduced in the cell monolayer. We next tested the influence of POSTN on cell invasion. HEYA8 cells were added to the top chamber of the Boyden chamber invasion assay and were allowed to migrate through pores to the opposite side of the membranes into either CM*^CTL^* or CM*^POSTNhigh^*, both of which contained 20% serum. Cells that had migrated through the membrane were then stained and counted. HEYA8 cells exhibited significantly more invasion into the CM*^POSTNhigh^* than the CM*^CTL^*environment, as shown in **Figure 2C** (p=0.006).

We next determined how POSTN influences response to carboplatin and paclitaxel, two commonly used chemotherapeutic agents for treatment of epithelial ovarian cancer. We used A2780 ovarian cancer cells cultured with either CM*^POSTNhigh^* or CM*^CTL^*. The CM*^POSTNhigh^* culture conditions enhanced chemoresistance to both drugs compared to cells grown with CM*^CTL^* conditions (**Figure 2D**) (p=0.049 and p=0.043 for carboplatin and paclitaxel, respectively). Results consistent with these findings were obtained using HEYA8 cells (*Supplementary data figure 1*).

### Periostin is protective against apoptosis

We next tested how elevated levels of extracellular POSTN influence ovarian cancer cell apoptosis. Apoptosis was induced by paclitaxel and followed by flow cytometry with propidium iodide (PI) staining for cell cycle analysis. As shown in **Figure 3A**, there were fewer apoptotic HEYA cells when they were cultured with CM*^POSTNhigh^* than when they were cultured with CM*^CTL^* (1.8% versus 13.9%, respectively). Similar results were observed for another ovarian cancer cell line, SKOV3, and using 1 mM staurosporine, another inducer of apoptosis (*supplementary data Figure 2*).

**Figure 3.**
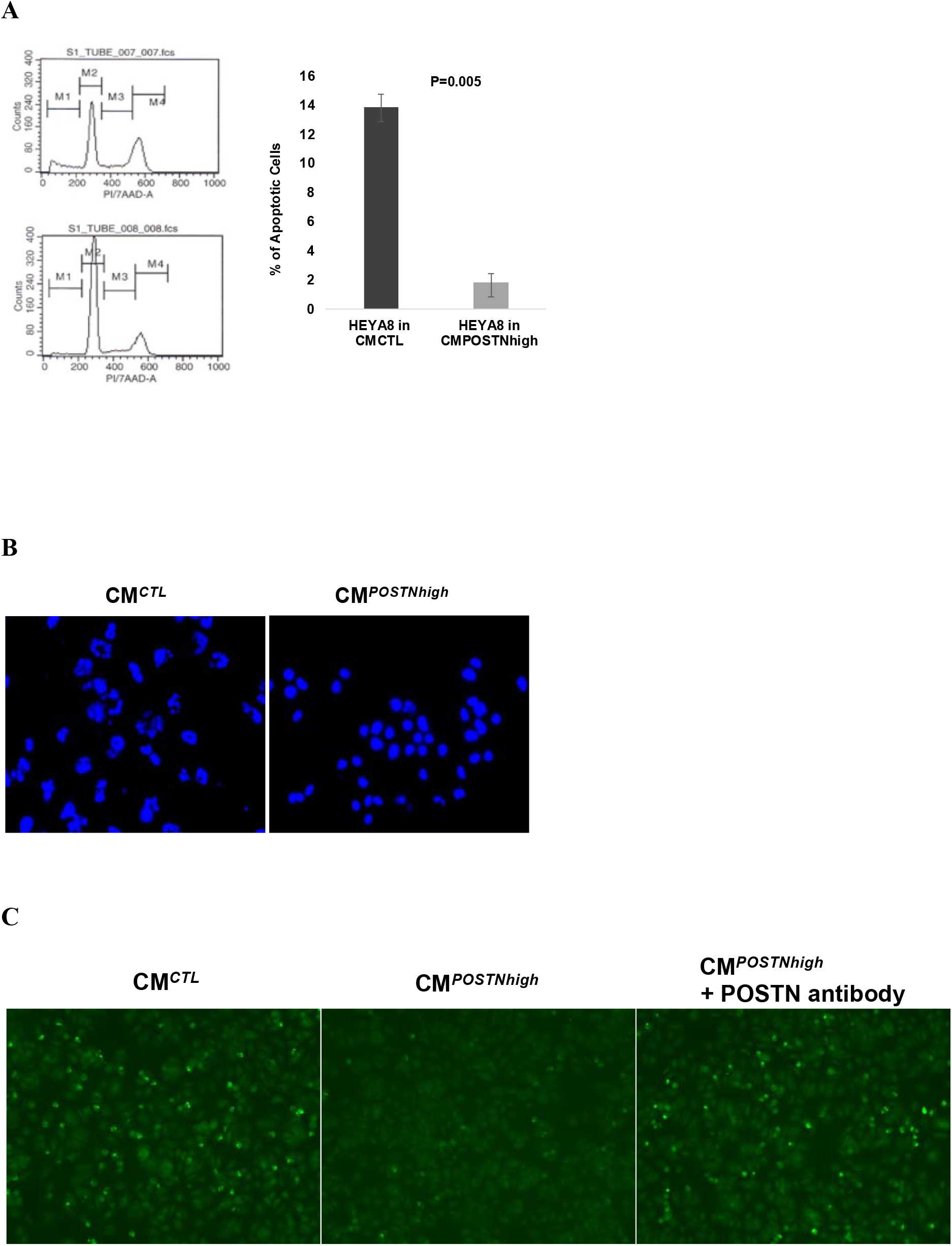
Increased resistance to apoptosis with high levels of POSTN in the microenvironment. **A.** HEYA8 cells exhibit a reduced number of cells undergoing apoptosis induced by paclitaxel (PTX) and cultured under CM*^POSTNhigh^* compared to cells grown under CM*^CTL^*. **B.** HEYA8 cell nuclei exhibit less fragmentation when treated with paclitaxel and cultured under CM*^POSTNhigh^*as compared to CM*^CTL^* conditions. The image was taken using a florescent microscope (20×10 magnification). **C.** Protection from apoptosis was reduced in HEYA8 cells under CM*^POSTNhigh^* when they were treated with an anti-POSTN antibody (10x magnification).

Nuclear fragmentation is a prominent morphological feature of cells undergoing apoptosis ^25^, so we examined cell morphology following staining with DNA content dye Hoechst 33342. The nuclei showed less fragmentation in the HEYA8 cells cultured with CM*^POSTNhigh^* compared to the cells cultured with CM*^CTL^* (**Figure 3B**), suggesting that extracellular POSTN contributes to apoptotic resistance. Caspases are critical mediators of apoptosis, and caspase-3 is a death protease frequently activated during this process ^26^. We therefore examined caspase-3 levels in HEYA8 cells cultured for 96 hours with either CM*^CTL^*, CM*^POSTNhigh^*, or CM*^POSTNhigh^* plus a monoclonal antibody to human POSTN using NucView® 488 caspase-3 assay kit (**Figure 3C**). Apoptosis was induced using 100 nM of paclitaxel for 24 hours. As shown in **Figure 3C**, caspase-3 activity was lower in HEYA8 cells when they were cultured with CM*^POSTNhigh^* (**Figure 3C** middle panel) than when HEYA8 cells were cultured with CM*^CTL^* (**Figure 3C** left panel). Inclusion of the neutralizing POSTN antibody with the CM*^POSTNhigh^* resulted in increased apoptosis, evident from the increase in caspase-3 activation (**Figure 3C** right panel), indicating that the apoptotic inhibition observed in cells cultured with CM*^POSTNhigh^* was alleviated by the POSTN-specific antibody. These data indicate that apoptosis inhibition in HEYA8 cells is partially attributable to specific effects of POSTN in the extracellular environment.

### POSTN-CM enhances the OC side population of stem-like cancer cells

Cancer stem cells have been shown in numerous cancer models to be involved in tumor development, cell proliferation, metastasis, and tumor recurrence due to their capacity for sustained self-renewal and genomic instability ^27, 28^. Several techniques have been developed to identify cancer stem cells including measurement, by FACS analysis, of the proportion of cells in a population that are able to efflux the Hoecsht dye H33342, referred to as a “side population” ^29^. To determine the stemness of OC cells regarding POSTN exposure, we performed a side population analysis. HEYA8 ovarian cancer cells were cultured in 3D culture conditions for 72 hours with either CM*^CTL^* or CM*^POSTNhigh^* followed by side population assay. The mean percentage of side population from the cancer cells cultured in CM*^POSTNhigh^* or CM*^CTL^* was calculated from two replicated tests, and the student t-test was used for the comparison between groups. The cells cultured with CM*^POSTNhigh^* exhibited a significantly larger side population compared to the control cells (p=0.027) (**Figure 4**). Similar results were obtained using CAOV2 and SKOV3 OC cell lines (*supplementary data Figure 3*).

**Figure 4.**
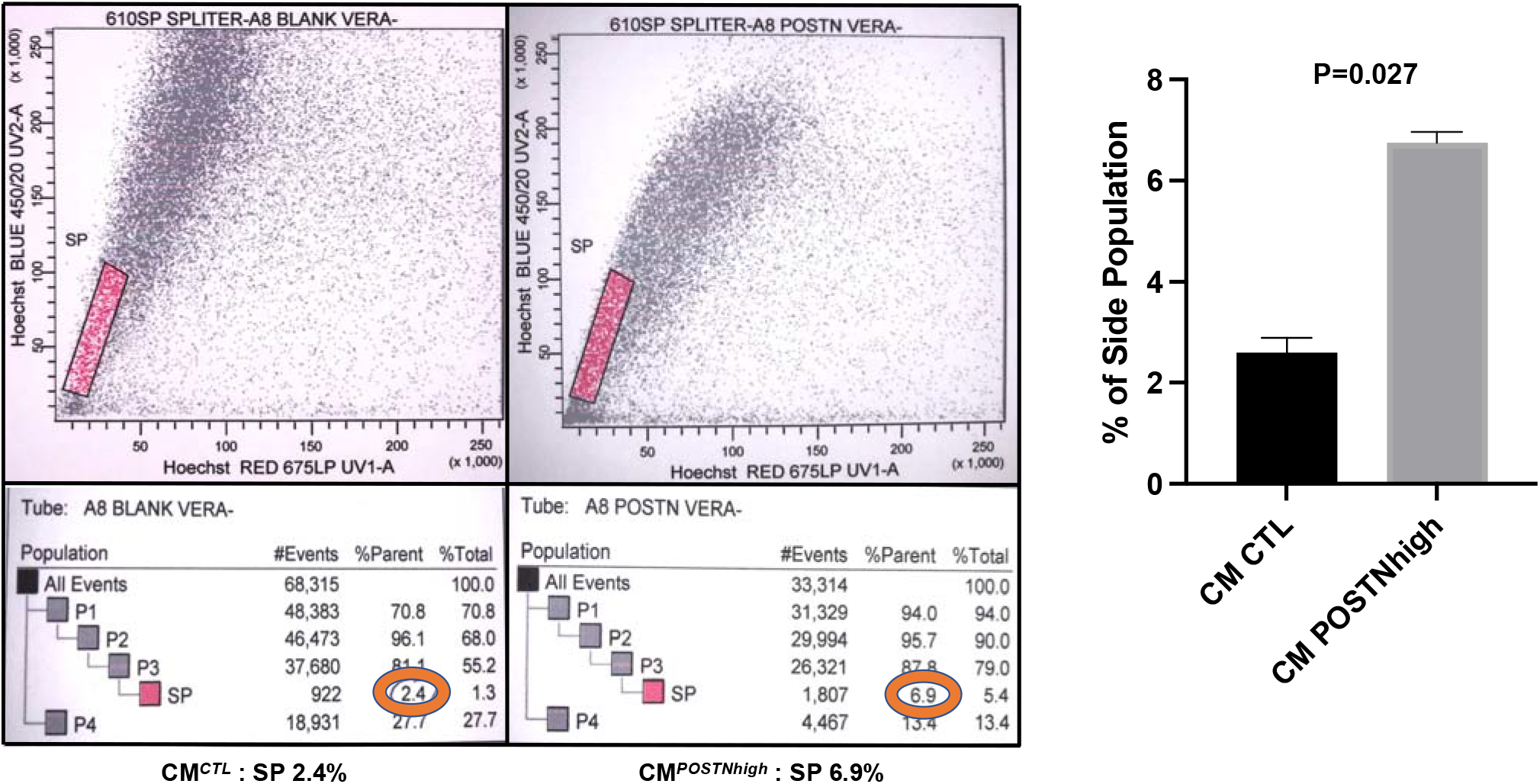
CM*^POSTNhigh^* culture enhances cancer cell stemness. The HEYA8 cancer stem cell-like side population (SP) increased in cells grown under CM*^POSTNhigh^* when assessed relative to CM*^CTL^*following H33342 fluorescence staining and verapamil treatment as a control. The average of side population from two independent tests with HEYA8 cells were 2.6% in CM*^CTL^* and 6.67% in CM*^POSTNhigh^* (p=0.027).

### Functions of POSTN in lipid metabolism in the cancer microenvironment and in cancer cells

When *POSTN* was overexpressed in 3T3-L1 preadipocyte cells (3T3-POSTN) via transfection with POSTN plasmid, we noticed cell morphology changed, and the cells were larger after cell differentiation was induced when compared to the 3T3-CTL cells. We then performed staining with Oil Red O which is a fat-soluble dye that stains neutral triglycerides and lipids. The 3T3-*POSTN* cells exhibited a higher proportion of lipids in the cytoplasm than did the 3T3-CTL cells (**Figure 5A**). This data suggests that *POSTN* overexpression in preadipocytes enhances lipid production and storage.

**Figure 5.**
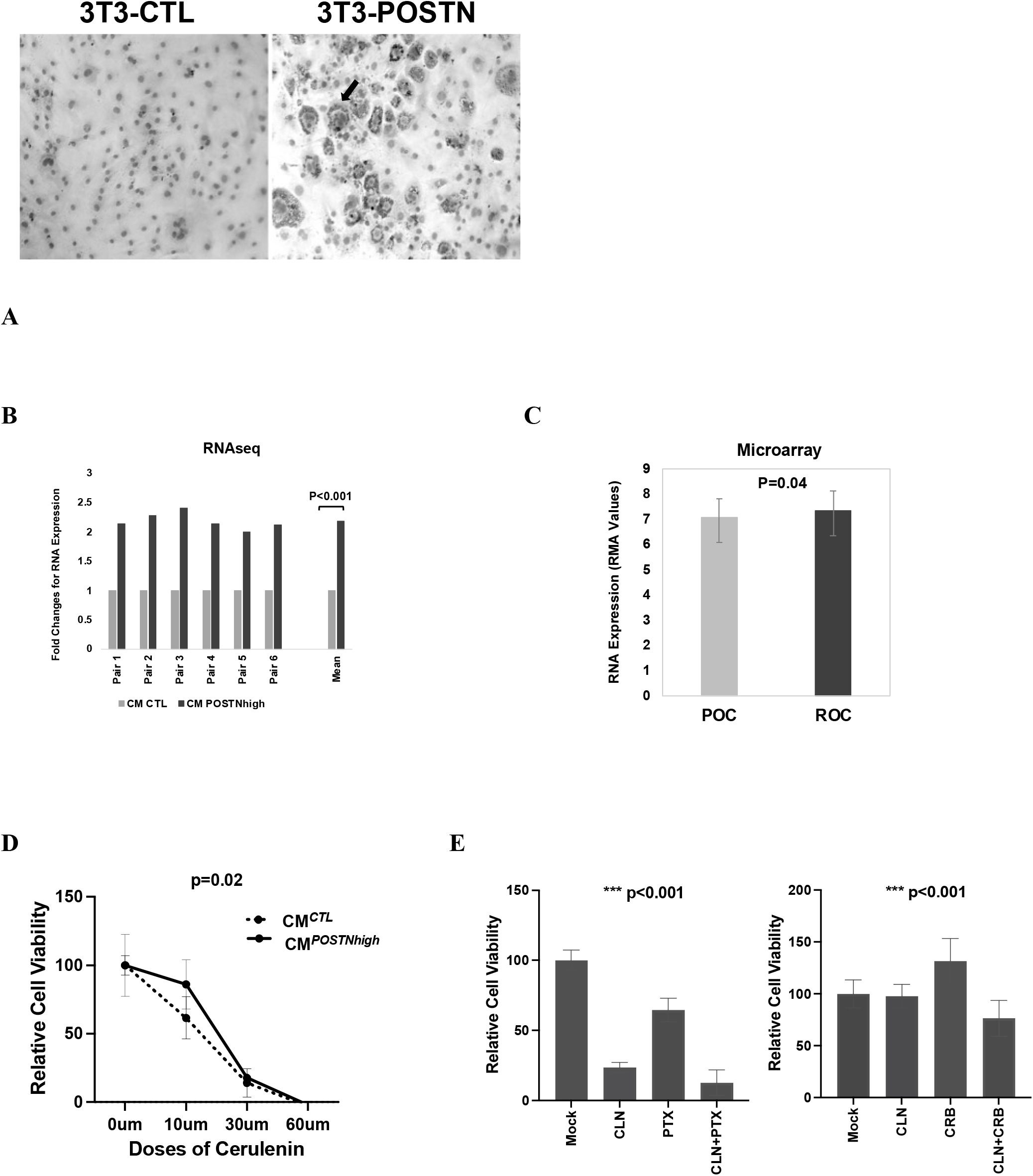
POSTN promoted lipid accumulation and is involved in lipid metabolism in the cancer microenvironment and in cancer cells. **A.** *POSTN-*overexpressing 3T3-L1 preadipocytes exhibit enhanced lipid production. Imaging was performed in differentiated cells after oil red O staining at 10×10 magnification. The lipid drops stained with oil red O was pointed by an arrow. **B.** RNA sequencing data shows that the fatty acid synthesis gene (*FASN*) is more highly expressed in HEYA8 cells cultured under CM*^POSTNhigh^*as compared to CM*^CTL^* (six replicates; p<0.001). **C.** *FASN* expression is higher in rOC that in matched pOC (p=0.04). **D.** HEYA8 cells under CM*^POSTNhigh^* were more resistant to FASN inhibitor Cerulenin (0-60 µM) than those under CM*^CTL^*(F=7.42, p=0.01). **E.** FASN inhibition combined with chemotherapy exhibits enhanced efficacy than either alone in HEYA8 cells under CM*^POSTNhigh^*. Left, cells treated with Cerulenin (CLN), Paclitaxel (PTX), or CLN+PTX; right, cells treated with CLN, Carboplatin (CRB), or CLN+CRB. The cancer cells were more sensitive to combined treatment as compared to single drug treatment (p<0.001 for CRB+CLN combined or PTX+CLN combined).

To explore the functional contribution of a lipid-enriched environment on cancer cell lipid metabolism, HEYA8 cells were cultured in 3D conditions with either CM*^CTL^* or CM*^POSTNhigh^* followed by RNAseq analysis (**Figure 5B**, left panel). We found expression of Fatty Acid Synthase (*FASN*) was significantly increased in cancer cells when cultured using CM*^POSTNhigh^*in all 6 repeated pairs (p<0.001). Additionally, we also found that the expression of *FASN* in recurrent OCs was significantly higher than in the matched primary OC (p<0.05) (**Figure 5C**). Our data suggested that FASN, which is important for catalyzing fatty acid synthesis in cancer, is functionally responding to the elevated levels of POSTN in the tumor microenvironment, the increased expression of FASN in cancer cells can lead to increased fatty acid synthesis. HEYA8 cells grown in CM*^CTL^*or CM*^POSTNhigh^* were treated with FASN inhibitor cerulenin (CLN). The cells showed more sensitivity to the FASN inhibitor when cultured in CM*^CTL^* than when cultured in CM*^POSTNhigh^* (**Figure 5D**, p=0.01). These results indicate that FASN functions in cancer cells is regulated by the levels of POSTN in the cancer microenvironment.

It has been shown the cerulenin can enhance antitumor activity when combined with chemotherapy reagent in human colon cancers ^30^. We therefore tested the effect of FASN on OC cells’ response to paclitaxel. HEYA8 cells grown in CM*^POSTNhigh^*medium were mock treated or treated with paclitaxel (PTX), cerulenin (CLN), or CLN+PTX (**Figure 5E**, left panel). The data show that CLN and PTX had an augmented effect on cell viability, and the effect was significantly stronger than the effect of either agent on its own (compared to PTX only: p < 0.0001, 51.9% decrease; compared to CLN: p = 0.0103, 11% decrease). This indicates that FASN inhibition modulates the effects of taxane-based chemotherapy on ovarian cancer cells. The same effect was also seen with a platinum-based chemotherapy agent, carboplatin (CRB) (**Figure 5E, right panel**). Using the single treatment, HEYA8 was resistant to CRB (p < 0.0001, 32% increase) and sensitive to cerulenin (p < 0.0001, 29% decrease). When these two reagents were combined (CLN+CRB), the chemoresistance effect observed with carboplatin alone (CRB) was reversed. This data suggests that when ovarian cancer exposed to a *POSTN*-high condition (high fat environment), the inhibition of FASN activity in cancer cells can enhance chemotherapeutic effects.

### Periostin is functionally regulated by Akt pathway

Studies have shown that the PI3K/AKT/mTOR pathway is activated in approximately 70% of ovarian cancer and that it plays roles in promoting cancer cellular growth, proliferation, and cell survival through an intricate series of hyperactive signaling cascades ^31^. To investigate the roles of the PI3K/AKT/mTOR pathway in cancer cells when exposed to a high POSTN environment, we cultured HEYA8 cells in 3D with either CM*^POSTNhigh^* or CM*^CTL^* medium, CM*^CTL^* + MK2206 (Akt inhibitor), or CM*^POSTNhigh^* + MK2206 for 72 hours. As shown in **Figure 6A**, the cancer cells formed spheroids in both CM*^CTL^*and CM*^POSTNhigh^* culture conditions. We were able to see a tighter and more spheroid formation for HEYA8 cells in CM*^POSTNhigh^*culture. The spheroids became smaller and disrupted when cells were exposed to MK-2206 in CM*^CTL^* medium. With MK-2206 present in CM*^POSTNhigh^*medium, there were even fewer spheroids, and they were more disrupted. Western blotting demonstrated that phospho-Akt was present at higher levels in the HEYA8 cells cultured with CM*^POSTNhigh^* compared to CM*^CTL^*-cultured cells (**Figure 6B**). The phospho-Akt was inhibited by MK-2206 in both POSTN-low (CM*^CTL^*) and POSTN-high (CM*^POSTNhigh^*) culture conditions.

**Figure 6.**
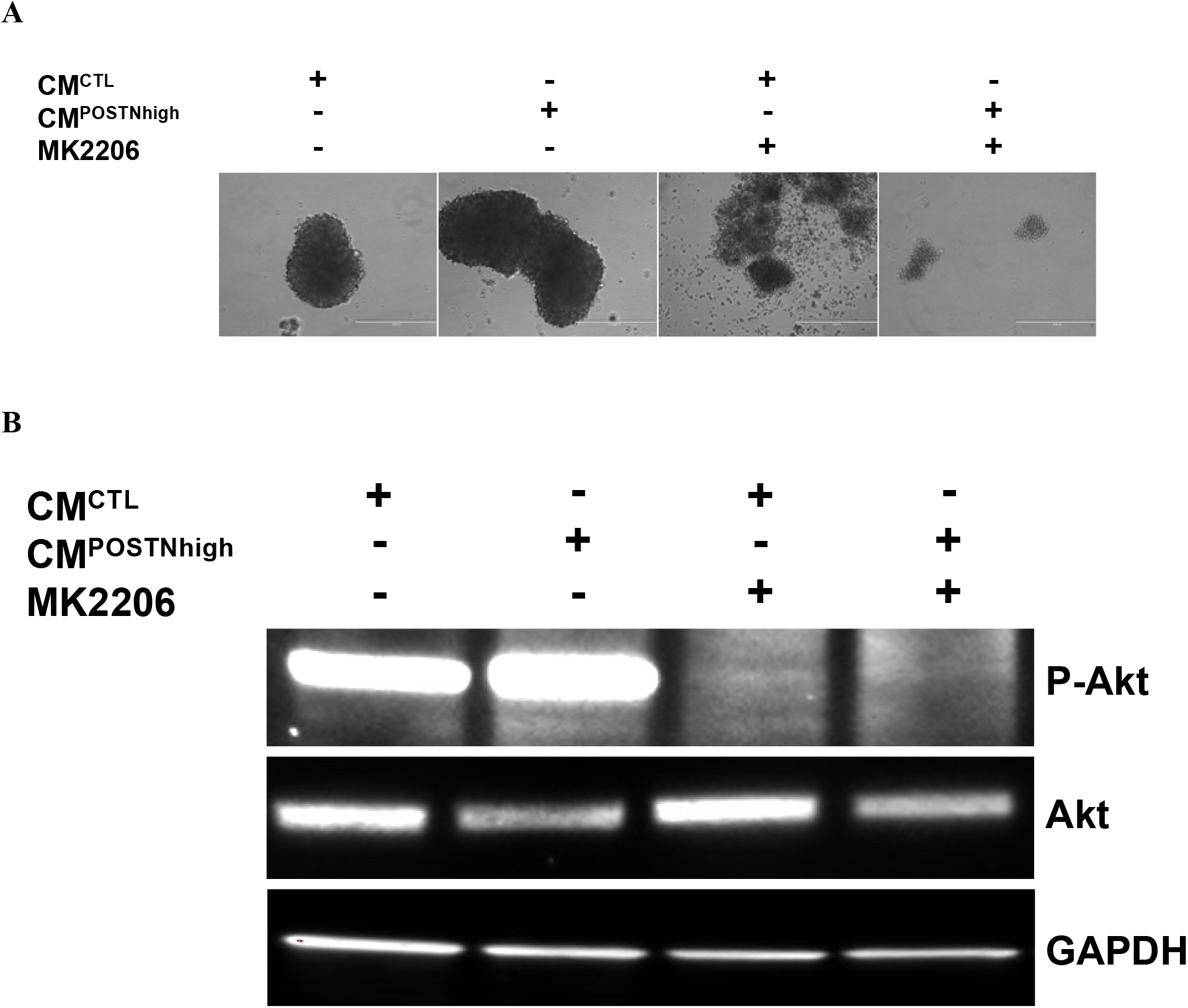
The Akt pathway contributes to regulating POSTN in cancer microenvironment. **A.** HEYA8 spheroid formation was disrupted under CM*^CTL^* and CM*^POSTNhigh^*conditions when cells were treated with Akt inhibitor MK-2206 with a more obvious disruption when the cells in . **B.** Akt activation is increased in HEYA8 cells when treated with Akt inhibitor MK-2206 under CM*^POSTNhigh^* as compared to CM*^CTL^*. Western blots are shown using an anti-phospho-Akt antibody (top panel) or anti-total Akt antibody (middle panel). GAPDH was used as the internal loading control (bottom panel).

### High levels of POSTN in the tumor environment enhances tumor formation in vivo

We used a xenograft mouse model of serous epithelial OC to determine if exogenous POSTN contributes to tumor formation *in vivo*. Two groups of female athymic nude mice were used, with 10 mice in the CM*^POSTNhigh^* group and 8 mice in the CM*^CTL^* group. On day 0, the mice in the CM*^POSTNhigh^*group were subcutaneously injected with CAOV2 cells (5x10^6^ per mouse) resuspended in CM*^POSTNhigh^* medium and ECM Matrigel at a 1:1 ratio. Mice in the CM*^CTL^* group were subcutaneously injected with the same number of CAOV2 cells resuspended in CM*^CTL^*medium and ECM Matrigel at a 1:1 ratio. Tumor length and width were measured on Days 11 and 18 after cancer cell injection using a caliper. On Day 20, the mice were euthanized, and tumor length and width were measured again after tumor removal (**Figure 7**). At Day 20, 9 out of 10 mice (90%) in the CM*^POSTNhigh^*group and 8 out of 8 mice (100%) in the CM*^CTL^* group showed tumor formation. Average tumor volume in the CM*^POSTNhigh^* group was larger, although not significant, on Day 11 (32.1 mm^3^ in CM*^POSTNhigh^* group vs 25.9 mm^3^ in CM*^CTL^*group, p=0.513, *supplementary data Figure 4)* and Day 18 (122.3 mm^3^ in CM*^POSTNhigh^* group vs 97.6 mm^3^ in CM*^CTL^* group, p=0.898, *supplementary data Figure 4)*. The difference was significant by Day 20 (228.4 mm^3^ in CM*^POSTNhigh^*group vs 144.6 mm^3^ in CM*^CTL^* group, p=0.0023; **Figure 7**).

**Figure 7.**
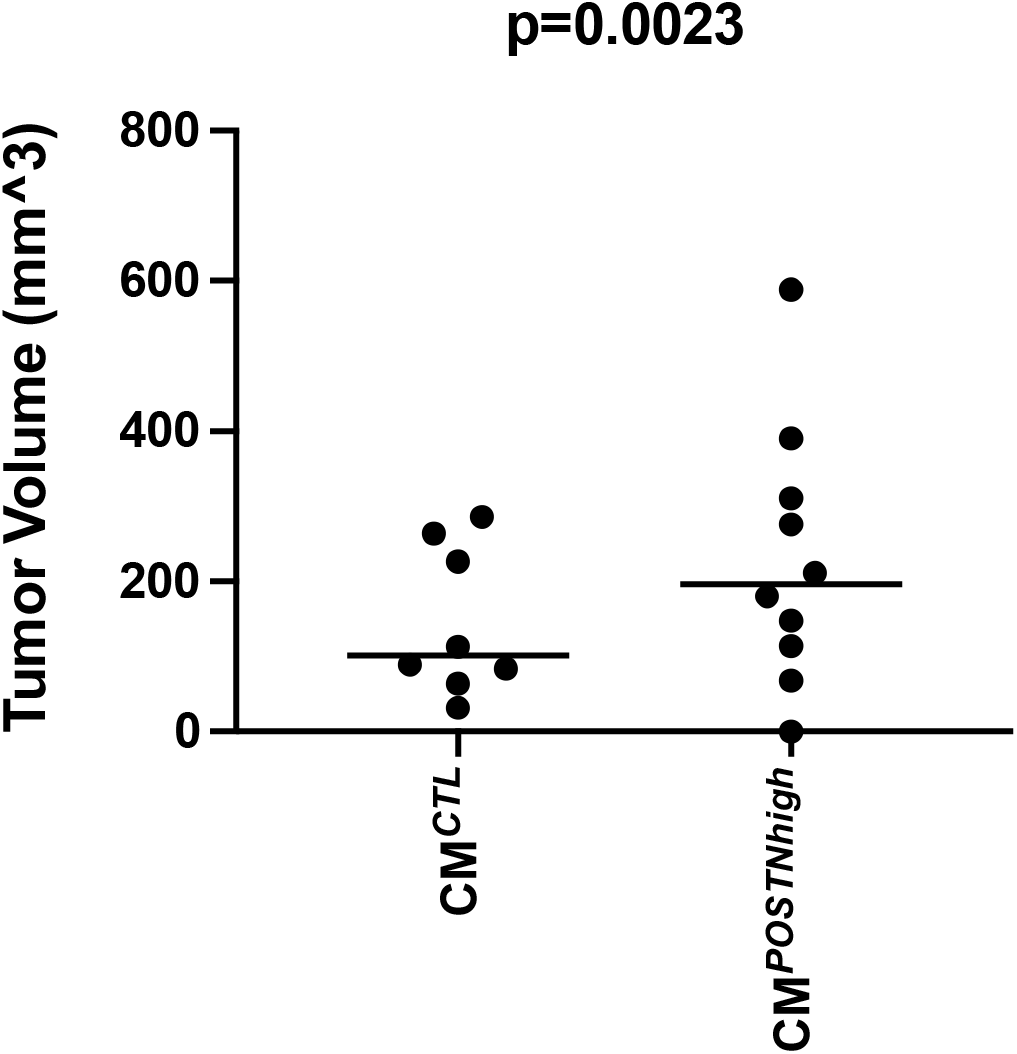
Ovarian cancer cells under CM*^POSTNhigh^* exhibit increased cancer formation *in vivo*. Female athymic nude mice were subcutaneously injected with 5x10^6^ CAOV2 cells suspended in CM*^POSTNhigh^* and ECM Matrigel at a 1:1 ratio (n=10). Mice in the CM*^CTL^* group (n=8) were injected with 5x10^6^ CAOV2 cells suspended in CM*^CTL^* and ECM Matrigel at a 1:1 ratio. Tumors were measured and tumor volume calculated on Day 20, after removal. Mean tumor volume was higher for cancer cells injected with CM*^POSTNhigh^* (mean= 228.4 mm^3^, 95% CI, 105.7 to 351.2) compared with CM*^CTL^*(mean= 144.6 mm^3^, 95% CI, 62.33 to 227.0) (p=0.0023).

## Discussion

In the present study, we report that *POSTN* is expressed in stromal tissues and by cancer cells and that expression is increased in recurrent ovarian cancer compared to primary ovarian cancers. We also found that the overexpression of *POSTN* gene correlates to short survival terms. We also found that higher POSTN levels in the TME contribute to a more aggressive phenotype, including increased proliferation and invasiveness, resistance to apoptosis, enhanced stemness, and increased chemoresistance, and contributed to increased tumor formation *in vivo*. More importantly, *POSTN* expression in preadipocytes (fibroblasts) led to increased lipid production and induced the expression inside cancer cells of the *de novo* fatty acid synthesis protein, FASN. These results indicate that lipid metabolism in stromal tissue is linked to lipid metabolism inside cancer cells.

We found that *POSTN* exhibits higher expression in recurrent relative to primary ovarian cancers. As POSTN has been detected in plasma ^32, 33^, our findings suggest that POSTN has the potential to serve as a diagnostic biomarker for rOC. In addition, the higher levels of *POSTN* expression, along with the phenotypes we identified that are associated with that higher level of expression, provide a foundation for future studies focused on targeted therapeutics for treatment of recurrent ovarian cancer.

Cancer stem cells (CSCs) are thought to be the source of recurrent disease ^34^. Enrichment of CSCs and their role have been intensively studied in cancer, including ovarian cancer recurrence ^35–37^. CSCs typically represent a small proportion of the total number of tumor cells, but they can drive both tumor development and progression. CSCs are not only responsible for primary tumor growth and metastasis, but also contribute to treatment failure, tumor progression, and relapse, since these cells are able to circumvent standard therapeutic approaches ^34^. As the elimination of this cell population is critical for increasing treatment success, a deeper understanding of ovarian CSC pathobiology is needed, including their niche; signals for differentiation that trigger recurrence, including epithelial-mesenchymal transition (EMT) ^38^; their signaling pathways; and how they interact with the tumor microenvironment. We found that a POSTN-high environment increased cancer stemness in ovarian cancer, evidenced by an increase in the CSC side population, suggesting the importance of TME factors in CSC behavior. Targeting the factors that enhance CSC numbers, such as POSTN in the TME, could provide a mechanism for reversing therapy resistance and reducing cancer recurrence.

We have shown that high expression of *POSTN* in cancer cell conditioned medium is associated with an increase in phospho-Akt levels, and that this is reduced by the addition of the Akt inhibitor MK-2206 (**Figure 6B**). Thus, there appears to be a link between the phosphorylation of Akt and higher levels of exogenous POSTN. Akt phosphorylation is frequently detected in ovarian cancer and can be targeted to disrupt ovarian tumor cell growth ^39–42^. Activation of the Akt signaling pathway increases the expression of apoptosis inhibitor protein survivin, allowing for improved cancer cell survival ^43^. Aberrant activity of Akt pathways in cancer is emerging as a focus for development of new cancer drugs ^44^. Our data showing the connection between *POSTN*-high expression in the TME in ovarian cancer and Akt activity support that targeting the Akt pathway could be a helpful strategy to treat OC patients with higher levels of *POSTN* expression.

We are the first to show that upregulation of *POSTN* in 3T3-L1 preadipocytes enhanced their lipid production (**Figure 5A**). This suggested that another role of POSTN in the TME is to support production of a lipid-based energy source and to possibly regulate lipid metabolism in cancer cells. Indeed, using RNAseq analysis, we were able to see higher expression of *FASN* in cancer cells cultured in POSTN-high conditioned medium. As a central regulator of lipid metabolism, FASN has been found to play a critical role in the growth and survival of tumors with lipogenic phenotypes^45^. Due to the role of FASN in *de novo* fatty acid synthesis, our data suggest that lipid metabolism in cancer cells may change through *FASN* overexpression when the cells are exposed to an environment with higher POSTN levels. Our data also provide evidence for a communication between fatty acid metabolism inside and outside of cancer cells. A recent study suggested that FASN inhibition increases mitochondrial priming and enhances breast cancer cell sensitivity to BCL2-targeting BH3 mimetics which may directly activate the apoptotic machinery and generate more potent and longer-lasting antitumor responses in a clinical setting ^46^. We showed that OC cells are more resistant to the FASN inhibitor Cerulenin when they are cultured in the conditioned medium with higher POSTN. The combination of FASN inhibition and chemotherapy reagents (carboplatin and paclitaxel) showed a cooperative effect in OC cells with POSTN-high media. These findings together suggest that a critical network between lipid metabolism in cancer cells and the TME may provide new inroads for OC treatment.

POSTN, produced by adipocytes or fibroblasts and secreted into the cancer extracellular matrix, can enhance cancer stemness. This, in turn, results in cancer cell chemoresistance which may contribute to cancer recurrence. Additionally, FASN induced in cancer cells by elevated exogenous POSTN plays important roles in lipid metabolism to generate excessive amounts of free fatty acids, which are then broken down into acetyl-CoA ^47^. Acetyl-CoA then supports mitochondrial respiration through fatty acid oxidation (FAO) to generate a greater amount of energy per unit mass compared to glucose ^48^. This could at least partly, if not completely, support cancer cell regrowth or recurrence.

Previous studies suggested that *POSTN* expression in OC cells promoted intraperitoneal tumor metastatic growth in immunodeficient mice ^49^. Using a neutralizing antibody to Periostin inhibited ovarian tumor growth and metastasis using a xenograft mouse model ^50^. Due to the preferential expression of POSTN in the stroma associated with cancer cells and the important connections between stromal POSTN and patient prognosis ^51^, we performed an *in vivo* study using CAOV2 OC cancer cells in the presence of POSTN-high CM. We found that tumors from OC cancer cells exposed to POSTN-high CM were significantly larger than tumors from OC cells in control CM.

In conclusion, we found that *POSTN* is significantly overexpressed in rOC compared with matched pOC. We also showed enhanced cancer stemness and lipid production when POSTN levels are elevated in the culture media, mimicking the TME. Our findings highlight the importance of POSTN in cancer stromal tissues in OC recurrence. Future studies will focus on how POSTN contributes to OC recurrence using patient-derived Xenograft models.

## Materials and Methods

### Tumor tissues

Primary and matched recurrent OC tissue sets (n=16 sets) were from patients with stage III/IV high grade serous epithelial ovarian cancer. The primary tumor specimens were collected at the time of initial debulking surgery. Recurrent tumor samples were obtained from the same patients during “second-look” surgeries. Samples were obtained after patients provided written informed consent and were stored in the Duke Gynecologic Oncology Tissue Bank under protocols approved by the Duke University Institutional Review Board.

### Microarrays

RNA was extracted from the paired primary-recurrent frozen tumor samples using the RNA Mini Kit according to the manufacturer’s protocol (Qiagen; Germantown, MD). Nucleic acid concentration and purity were assessed using a NanoDrop^TM^ 2000 spectrophotometer (Thermo Fisher Scientific; Waltham, MA). RNA (1 µg) was analyzed for expression using the Affymetrix Human Genome U133 Plus 2.0 microarray, which included 22,277 probes. The resulting gene expression data were normalized using the robust multiarray average algorithm (RMA) ^52^. The Affymetrix gene expression data were analyzed by comparing values for primary and recurrent tumors using a paired t-test (alpha = 0.05), adjusted for multiple testing using the Bonferroni correction.

### Immunohistochemistry

Frozen tumor tissues were cut into 5 _μ_m sections using a microtome and the sections placed on slides. Immunohistochemistry staining for cytokeratin and POSTN was carried out according to a previously published protocol ^53^. Briefly, after blocking non-specific binding using blocking buffer, normal goat serum (Abcam, Ab7481), the slides were incubated at 4°C overnight with a monoclonal antibody to human cytokeratin antigen (Abcam, Cat #ab53280, at 1:250), or monoclonal antibody to human POSTN (R&D Systems, Cat # AF3548, at 10 µg/mL), or anti-human IgG (Abcam, Cat#109489, at 1:500). Immunodetection was carried out using the streptavidin-biotin-based Multi-Link Super Sensitive Detection System (4plus Universal HRP Detection System, Biocare Medical). Immunodetection was performed using an ZEISS light microscope. Micrographs were taken at 10×10 magnification.

### Cell culture

Mouse 3T3L1 fibroblast cells were maintained in Dulbecco’s Modified Eagle Medium with high glucose (MilliporeSigma, Cat # D5796), 10% Bovine Calf Serum (MilliporeSigma, Cat # 12133C) and 1% Penicillin-Streptomycin (MilliporeSigma, Cat #p4333). HEK293T cells were maintained in DMEM with high glucose, 10% Fetal Bovine Serum (FBS, ThermoFisher, Cat # 10082147) and 1% Penicillin-Streptomycin. All ovarian cancer cell lines were maintained in RPMI 1640 (ThermoFisher, Cat # A4192301) with 10% FBS and 1% Penicillin-Streptomycin. Cells were incubated at 37°C in a humidified chamber with 5% CO_2_. Human cells were genetically authenticated by the Duke University DNA Analysis Facility and confirmed to be free of mycoplasma by the Duke Cell Culture Facility.

### Generation of POSTN conditioned medium

3T3-L1 cells were stably transfected with a human *POSTN* lentiviral vector (pLenti-GIII-CMV-RFP-2A-Puro-*POSTN*, Applied Biological Materials Inc. Cat# LV268492) and control vehicle plasmid (pLenti-CMV-RFP-2A-Puro-Blank Vector, Applied Biological Materials Inc. Cat# LV591) to generate 3T3-L1 cell lines that do (3T3-*POSTN*) and do not (3T3-CTL) overexpress *POSTN*, respectively. Lentivirus transfection was carried out according to the protocol described previously ^54^. Due to the co-expression of red fluorescent protein (RFP) in cells, we were able to select *POSTN*-positive cells using flow-activated cell sorting. Overexpression of *POSTN* in 3T3-L1 cells (3T3-*POSTN*) as compared to 3T3-CTL cells was confirmed using RT-PCR with a human *POSTN*-specific probe as described below. POSTN protein expression in the medium was confirmed using an ELISA assay with an anti-POSTN polyclonal antibody (R&D Systems, Cat # AF3548). 3T3-CTL cells were used to generate “control” conditioned medium (CM*^CTL^*), and 3T3-*POSTN* cells were used to generate conditioned medium with higher POSTN levels (CM*^POSTNhigh^*). Media were collected for use after 48-72 hours of culture and centrifuged at 1500 g for 5 minutes to remove cellular debris. This conditioned medium (CM) was used for OC cell culture experiments as described and is referred to as CMCTL or CMPOSTNhigh.

### RT-PCR

Total RNA was extracted using RNA STAT-60 reagent according to the manufacturer’s instructions (AmsBio, Cat # CS-110). RT-PCR was carried out using 500 ng of total RNA in a 20 µL volume using SuperScript IV One-Step RT-PCR kit according to the manufacturer’s protocol (ThermoFisher, Cat# 12594025) with a Taqman probe specific to *POSTN* (Hs01566750, ThermoFisher, Cat# 4331182). Human Beta-2-Microglobulin (B2M) (ThermoFisher, Cat# 4333766T) served as an endogenous control for RNA input. The PCR reaction was performed at 95°C for 10 minutes followed by 40 cycles of 95°C for 15 seconds and 60°C for 1 minute. Relative RNA expression values were calculated using the threshold cycle (CT) values. This test was repeated three times independently with 6 replicates for each condition and each test.

### ELISA

Enzyme-linked immunosorbent assays (ELISA) using the human POSTN ELISA kit from Aviva Systems (Cat # OKCD09048) and the antibody included in the kit were performed according to the manufacturer’s instructions to compare POSTN protein expression in the CM*^CTL^* versus CM*^POSTNhigh^*. 3T3-CTL and 3T3-POSTN cells were cultured for 48-72 hours followed by the collection of medium. After brief centrifugation to remove cellular debris, the CM was collected and used for ELISAs. Total protein concentration was determined using Pierce™ BCA Protein Assay Kit (ThermoFisher, Cat# 23225). The total protein concentration was determined using Bradford protein assay according to the instructions from the manufacturer (Bio-Rad). POSTN protein expression from the ELISA assay was normalized with the total protein levels. A t-test was used to compare POSTN expression. This test was repeated two times independently.

### Wound healing

OC cancer cells were cultured in 5 mL of either CM*^CTL^* or CM*^POSTNhigh^* in 6-well plates. Following incubation or transfection for 24 hours, gaps were created in the cell monolayers using a sterile p200 pipet tip. Photomicrographs at 4x magnification were taken at indicated time points using a Zeiss inverted microscope.

### Invasion assay

A cell invasion assay kit (24-well; with a basement membrane) was purchased from Cell Biolabs (Cat # CBA-110). A HEYA8 OC cell suspension containing 150,000 cells was added to each insert of the invasion plate in 500 _μ_L of either CM*^CTL^* or CM*^POSTNhigh^*. CM*^CTL^* or CM*^POSTNhigh^*, each containing 20% FBS, was added to the lower chamber of the invasion plate. The cells were incubated for 48 hours at 37°C in a humidified chamber containing 5% CO_2_. Cells on the bottom of the invasion membrane were stained and quantified at OD 560 nm using a plate reader (BMG Labtech, POLARstar Omega) according to the protocol provided by the manufacturer (Cell Biolabs). This test was repeated three times independently.

### Chemosensitivity test

OC cell line A2780 was seeded into a 96-well plate (5,000 cells/well) in 100 μL of either CM*^CTL^* or CM*^POSTNhigh^* medium. After 24 hours, carboplatin or paclitaxel were added at the indicated dose ranges. Following 72 hours of drug treatment, cell viability was tested using the CellTiter-Glo® Luminescent Cell Viability Assay kit (Promega, Cat# G7570) according to the manufacturer’s instructions. The luminescence signal was recorded using a 96-well plate reader (BMG Labtech, POLARstar Omega). Readings were standardized to the mock treatment for each CM type and are reported as the percentage of viable cells relative to the mock treated cells. This test was repeated three times independently.

For chemosensitivity testing with cerulenin, HEYA8 cells were seeded into a 96-well plate at 4,000 cells/well with 100 μL of CM*^CTL^* or CM*^POSTNhigh^*. The cells were grown for 24 hours followed by treatment for 72 hours with either cerulenin at 10 μM, carboplatin at 3 μM, paclitaxel at 3 μM, 3 μM carboplatin + 10 μM cerulenin, or 3 μM paclitaxel + 10 μM cerulenin. At the end of the treatment, the CellTiter-Glo® Luminescent Cell Viability Assay kit reagent was added. Luminescence was measured using a 96-well plate reader. Readings were normalized to the control absorbance for each CM type, and relative to the mock treatment. This test was repeated three times independently.

### Cell apoptosis analysis by flow cytometry

HEYA8 cells were cultured for 96 hours with either CM*^CTL^*or CM*^POSTNhigh^*. Apoptosis was induced for 24 hours using paclitaxel at final concentrations of 100 nM with DMSO as a vehicle control. After incubation, cells were collected and resuspended in PBS/1%FBS. After fixation using ice-cold 70% EtOH for two hours, cells were washed using PBS/1%FBS. Propidium iodide (PI) at 50 µg/mL and RNase A at 10 µg/mL were added to cells. Apoptosis was analyzed with a flow cytometer using 488 nm excitation and emission collected at 575–610 nm. The cell cycle profile was obtained with the “sub-M1” peak representing the apoptotic population. This test was repeated three times independently.

### Cell apoptosis analysis by caspase-3 staining

HEYA8 cells were cultured for 96 hours with either CM*^CTL^*, CM*^POSTNhigh^*, or CM*^POSTNhigh^* + a mAb for human *POSTN* (R&D Systems, Cat # AF3548, at 10 µg/mL). Apoptosis was induced by treating cells with 100 nM paclitaxel for 24 hours. Apoptosis was analyzed using the NucView® 488 caspase-3 assay kit (Biotium, Cat# 10402-T) which contains NucView® 488 Caspase-3 substrate. Counterstaining was done using the Hoechst 33342 DNA dye included in the kit. The staining was carried out according to the protocol provided by the manufacturer. Images were obtained using a fluorescent microscope, AMG EVOS Imaging Microscope (AME-3206), at 20×10 magnification. This test was repeated two times independently.

### 3D cell culture

OC cells were cultured in 5 mL of either CM*^CTL^* or CM*^POSTNhigh^* in ultralow attachment 6-well plates (MilliporeSigma, Cat # CLS3471). Spheroid formation was visualized using an inverted microscope (Zeiss) after 72 hours of cell culture.

### Side population analysis

After 72 hours of 3D culture in either CM*^CTL^* or CM*^POSTNhigh^*, the side population of OC cells (HEYA8, CAOV2, and SKOV3) was measured using the flow cytometry approach ^29, 55, 56^. Briefly, after culture, the OC cells were collected and stained for 90 minutes at 37°C using 5 µg/mL Hoechst 33342 (bisBenzimide H 33342 trihydrochloride, MilliporeSigma, Cat# B2261). Verapamil (MilliporeSigma, Cat# V4629), an inhibitor of ABC transporters, was used as a negative control and was added to a final concentration of 50 μM along with the Hoechst 33342 dye. Before FACS analysis, propidium iodide (PI) solution was added to a final concentration of 1 μg/mL to identify nonviable cells. FACS analysis was carried out with a dual laser flow cytometer (Becton Dickinson FACS Vantage SE cell sorter). Hoechst dye excitation at 355-nm ultraviolet laser was measured at two wavelengths using a 424/44 (Hoechst blue) band-pass filter and a 585/42 (Hoechst red) band-pass filter. PI excitation was induced using a 488-nm laser and detected after passing through a 630/22 band-pass filter. PI–positive dead cells and debris were excluded. The side population from cancer cells cultured with CM*^CTL^* or CM*^POSTNhigh^*was compared using a paired student t-test. The experiments were repeated twice each with OC cell lines HEYA8, CAOV2, and SKOV3.

### 3T3-L1 differentiation and oil red O assay

To induce 3T3L1-CTL and 3T3L1-*POSTN* cells to differentiate into adipocyte-like cells, the cells were treated using the reagents from the 3T3-L1 differentiation kit according to instructions (MilliporeSigma, Cat #DIF001). Briefly, the cells were cultured until confluent with preadipocyte medium (DMEM medium with 10% bovine calf serum, 100 units/mL penicillin, and 100 μg/mL streptomycin). The preadipocyte medium was then replaced with differentiation medium containing 1 μL/mL of Differentiation Cocktail in DMEM/F12 (1:1) medium with 10% FBS. Three days after the differentiation, the medium was replaced with 1 μL/mL of insulin containing DMEM/F12 (1:1) with 10% FBS. Lipid droplet formation and accumulation was visible by light microscopy about 7-10 days after addition of the differentiation medium. The cells were fixed with 10% formalin and stained with oil red O (MilliporeSigma, Cat #MAK194) nine days after differentiation initiation. Micrographs were taken using an inverted microscope (Zeiss). This test was repeated three times independently.

### RNAseq

Due to low endogenous expression of *POSTN* based on microarray gene expression data for OC cell lines, HEYA8 cells were used in this study. HEYA8 cells were cultured in either CM*^CTL^* or CM*^POSTNhigh^* for 72 hours under 3D cell culture conditions. RNA was extracted using RNA STAT-60 (Amsbio, Cat#CS-111). We confirmed the increase in cell proliferation and invasion and apoptosis inhibition in the cells cultured in CM*^POSTNhigh^* before submitting 1 μg of RNA to the Genomic Analysis and Bioinformatics Core at Duke University for generation of RNA Seq data using the NovaSeq 6000 system. Differential gene expression analyses were performed by the same core to determine that the genes are expressed at different levels between the CM conditions.

### Western blotting

HEYA8 cells were seeded at equal cell numbers (1x10^6^) onto 100 mm ultralow attachment Petri dishes for 3D culture as described above. Each cell line was plated in four conditions: CM*^CTL^*, CM*^POSTNhigh^*, CM*^CTL^*+ 10 μL MK-2206 2HCl (a selective AKT inhibitor from Selleckchem, Cat#S1078), and CM*^POSTNhigh^* + 10 μL MK-2206 2HCl. The Evos FL Cell Imaging system was used to take micrographs of 3D cell cultures every 24 hours to check spheroid formation. Cells were harvested after 72 hours of culture for protein analysis with western blotting. The cell pellets were incubated in NP-40 cell lysis buffer (ThermoFisher, Cat#85124) for 30 minutes on ice. Cell lysates (50 µg) were loaded into 4–20% PROTEAN® TGX™ Precast Protein Gels (BioRad, Cat# 5678081). After overnight transfer to nitrocellulose membranes, the blot was incubated with primary antibodies, including anti-phospho-Akt (1:2500, Cell Signaling, Cat# #9271), anti-Akt (1:1000, Cell Signaling, Cat#9272), and anti-GADPH (1:2000, ThermoFisher, Cat# PA1-987). After secondary antibody incubation, ECL

Western Blot Substrate kit (Bio-Rad, Cat# 1705060S) was used to detect the signals, which were imaged using the Chemidoc imaging system from Bio-Rad.

### Ovarian cancer xenograft mouse model

Eighteen six-week-old female athymic nude mice were purchased from Charles River Laboratories. CAOV2 cells were prepared either in CM*^POSTNhigh^* or CM*^CTL^* medium and then mixed at a 1:1 ratio with Matrigel Growth Factor Reduced (GFR) Basement Membrane Matrix (Corning, Cat#354230). Ten mice were injected subcutaneously in the upper right flank with 5x10^6^ cells each of CAOV2 grown with CM*^POSTNhigh^*(CM*^POSTNhigh^* group). Eight mice were used as controls and were injected in the same manner with 5x10^6^ cells each of CAOV2 grown in CM*^CTL^* mixed with GFR Matrigel at a 1:1 ratio (CM*^CTL^*group). Tumors were surgically removed 20 days after cancer cell injection. Tumor length and width were measured using calipers and then the tumors were frozen at -80°C. Tumor volume was calculated using the formula 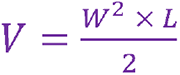57

### Statistical analysis

Chemosensitivity assays were evaluated using a one-way ANOVA test with replication. Gene expression from qRT-PCR was analyzed and compared between groups using Prism 9 (GraphPad Software, LLC). In mouse tumor study, the mean tumor volumes were compared between the CM*^CTL^* and CM*^POSTNhigh^* groups using the Mann Whitney U test. P values < 0.05 were considered statistically significant.

